# Interface-mediated secondary phase separation

**DOI:** 10.64898/2026.09.04.749388

**Authors:** Kaifeng Weng, Jie Lin

## Abstract

Inside living cells, many types of biomolecular condensates coexist and interact. Recent experiments have shown that new phases often form at the surface of preexisting condensates. However, the mechanisms for this interface-mediated phase transition remain elusive despite its importance to numerous biological processes. Here, we show that interface-mediated secondary phase separation is a universal pathway for forming a new phase in multicomponent solutions. Using Cahn-Hilliard simulations, we successfully generate puncta of client protein on the surface of scaffold condensates. Based on a quasistatic protocol, we theoretically demonstrate that an interfacial instability triggers new-phase formation and predict the amount of client protein needed for the instability to occur. Remarkably, our quasistatic theory successfully predicts the onset of secondary phase separation even if the client is rapidly added or both the client and scaffold are rapidly added. In the latter case, coarsening of the primary condensates drives secondary phase separation. Our work reveals that cells can exploit existing condensates to lower nucleation barriers of new phases, with important implications for protein aggregation.

## Introduction

Biomolecular condensates have emerged as a ubiquitous mechanism for spatial organization of the intracellular environment [1–5]. While most of our intuitive understanding of biomolecular condensates comes from liquid-liquid phase separation of binary mixtures [6–10], intracellular environments are complex with numerous components and multiple phases [11–14]. Thus, understanding the kinetic pathways of phase formation in a multicomponent system with more than two phases is critical.

Theoretically, even predicting the possible equilibrium states, including both metastable states and the ground state (i.e., the state with the lowest free energy), of a multicomponent solution is already a formidable challenge. Linear stability analysis (i.e., calculating the Hessian matrix based on the local free energy density) can inform us whether a uniform state is stable; however, it cannot tell us how many phases the equilibrium state has [15, 16]. Convex-hull analysis can tell us what the ground state is [17–19]; however, it cannot tell us whether the system can reach the ground state or arrest at other metastable states. Indeed, earlier simulations have shown that the equilibrium configurations for a ternary system can be more complex than the expected ground state [20, 21].

In biological systems, a new phase often forms in the presence of preexisting condensates. For example, in the experiments by Yan et al. [22], the authors found that stress granules (SG) promote the formation of TDP-43 aggregation localized on their surfaces *in vivo*. In the corresponding *in vitro* experiments, the authors added TDP-43 to a solution with preexisting SG-like condensates. They found that TDP-43 readily partitioned into these condensates. Strikingly, TDP-43 formed distinct puncta on the condensate surface after a finite time, mirroring intra-condensate demixing observed in cells. Apparently, the presence of preexisting condensates significantly reduces the energetic barrier to forming a new phase enriched in TDP-43, which cannot form spontaneously in the absence of preexisting condensates. However, the conditions for forming a new phase remain elusive, even in the simplified scenario of ternary mixtures.

In this paper, we tackle these critical issues by studying the kinetic pathways toward three-phase coexistence in a ternary mixture, which is simple enough but still highly representative of real systems [22]. We demonstrate that preexisting condensates significantly lower the energetic barriers for secondary phase separation of the client protein. Our numerical simulations based on Cahn-Hilliard dynamics successfully reproduce experimental observations. More importantly, we show that an interfacial instability mechanism triggers the secondary phase separation. To show this, we implement a protocol in which we add the client quasistatically, allowing us to predict the equilibrium profiles. As the client is quasistatically added, an increasing amount of client protein accumulates at the condensate surface and eventually triggers an interfacial instability, leading to the formation of a new phase.

Our quasistatic theory predicts the minimum amount of client protein needed to trigger interfacial instability. Strikingly, the quasistatic theory works even if the client is added rapidly or if the client and scaffold are added rapidly simultaneously. Remarkably, when the client and scaffold are added simultaneously, coarsening of scaffold condensates (i.e., primary condensates) triggers the formation of the new phase. Specifically, the coarsening process continuously reduces the interfacial areas of the primary condensates such that the surface density of client protein continuously increases, eventually triggering the interfacial instability. We also demonstrate that the secondary phase separation always first forms on the surface of the largest primary condensate.

These striking observations suggest that interface-mediated secondary phase separation is a universal pathway that generates new phases in multicomponent solutions, which has far-reaching implications for the regulation of condensate morphologies and even irreversible aggregate formation on condensate surfaces [22].

### Model

We consider an incompressible ternary liquid mixture comprising a scaffold protein (*ϕ*_1_), a client protein (*ϕ*_2_) and a solvent molecule (*ϕ*_3_). The local volume fractions satisfy the incompressibility constraint *ϕ*_3_ = 1 − *ϕ*_1_ − *ϕ*_2_. In this work, we also use *ϕ*_*i*_ to denote the name of the components if possible. The thermodynamics of the system are governed by the following free energy density that has a gradient penalty [23]:

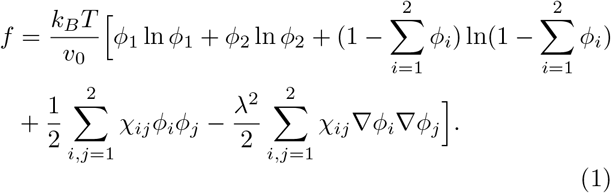

Here, *k*_*B*_ is the Boltzmann constant, *T* is the temperature, *v*_0_ is the molecular volume. *χ*_*ij*_ is the symmetric matrix that quantifies the interaction between component *i* and *j* and *λ* dictates the interfacial width. In this work, we consider the canonical ensemble where the average volume fractions over the whole space, 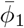 and 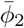, are fixed.

To derive the kinetic equations governing the spatio-temporal evolution of *ϕ*_*i*_, we employ the Onsager variational principle, assuming a frictional dissipation generated by relative motion between components (see detailed derivations in Supplemental Material (SM) section A1) [7]. The dynamics of the volume fractions, *ϕ*_1_ and *ϕ*_2_ satisfy the Cahn-Hilliard equations:

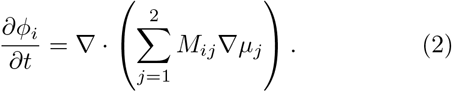

Here, the two-by-two mobility matrix, *M*_*ij*_ = *Dϕ*_*i*_(*δ*_*ij*_ − *ϕ*_*j*_), where *D* is the diffusion constant. The dimensionless exchange chemical potential satisfies

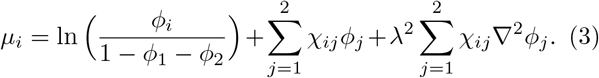

In the following simulations, the length unit is the system size *L* and the time unit is *λ*^2^*/D*. Details of numerical simulations are included in SM section A2-3, B1.

### Interface-mediated secondary phase separation

We first perform a two-dimensional simulation to test whether the simple model can capture the experiments in Ref. [22] where TDP-43 was added to a solution with preexisting SG-like condensates, which quickly got recruited into the condensates; after about 4 hours, TDP43 started to demix from the condensates and formed a new TDP-43-enriched phase on the surface of the preexisting condensates. To mimic the experiments, we initialize the system with a preexisting condensate enriched in *ϕ*_1_ in the center and then rapidly add *ϕ*_2_ at *t* = 0 (Fig. 1a; see SM Section B3 for protocol details). We introduce a mutual attraction (*χ*_12_ = −4.5) between *ϕ*_1_ and *ϕ*_2_ and set the *χ*_*ij*_ parameters so that the system’s ground state has three phases, thereby possessing the capacity for secondary phase separation.

**FIG. 1.**
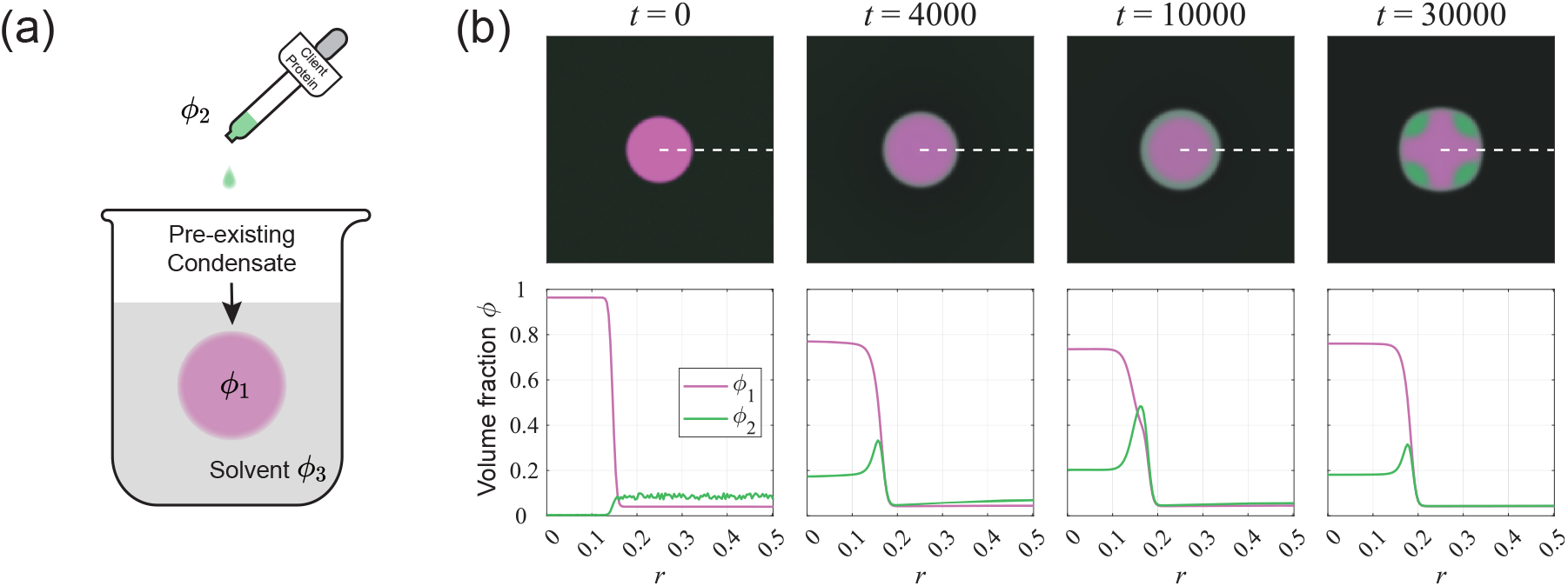
Interface-mediated secondary phase separation. (a) A common experimental protocol to induce three-phase coexistence in which a client protein is added to a solution with preexisting condensates of the scaffold protein. (b) Spatiotemporal evolution of the *ϕ*_1_ and *ϕ*_2_ fields (top) and corresponding one-dimensional profiles along the white dashed line (bottom). *ϕ*_2_ initially rapidly gets absorbed into the preexisting condensate, then gradually accumulates on the condensate surface, and finally exhibits a spontaneous symmetry breaking, generating a new phase enriched in *ϕ*_2_ that wets the preexisting condensate.

Interestingly, a three-stage kinetic pathway to the three-phase state emerges (Fig. 1b). In the first stage, *ϕ*_2_ is rapidly drawn from the solvent toward the condensate due to attraction between *ϕ*_1_ and *ϕ*_2_ (compare the snapshots at *t* = 0 and *t* = 4000). Interestingly, besides being enriched in the preexisting condensate, *ϕ*_2_ also preferentially accumulates at its surface. This leads to the second stage: an interfacial layer of *ϕ*_2_ around the condensate gradually accumulates (compare the snapshots at *t* = 4000 and *t* = 10000). Ultimately, the interfacial layer undergoes spontaneous symmetry breaking and a new phase enriched in *ϕ*_2_ appears, wetting the preexisting condensate. Our simulations nicely reproduce the phenomena observed in the experiments. We note the *ϕ*_2_ value inside the preexisting condensate only changes mildly before and after interfacial instability (Fig. 1b), in agreement with the experiments.

We seek to understand the quantitative conditions for the secondary phase separation to occur. In what follows, we consider a simpler scenario in which *ϕ*_2_ is added quasistatically, allowing us to theoretically predict the onset of interfacial instability. Later, we show that the quasistatic theory also applies to more complex scenarios. *Interfacial instability during quasistatic addition—*To map the boundaries between the two-phase and three-phase regions, we quasistatically increase the average volume fraction of *ϕ*_2_ (see SM Section B3 for details on the addition protocol) and monitor when interfacial instability occurs (Fig. 2a left). Remarkably, a preexisting condensate significantly lowers the instability threshold (the blue line in Fig. 2a left) compared with a solution without the preexisting condensate (i.e., the bulk spinodal threshold, which is the red dashed line in Fig. 2a left). We highlight a kink in the boundary curve around *χ*_12,*c*_ ≈ − 1. For *χ*_12_ ≳ − 1, the preexisting condensate acts as an excluded volume; in this region, the formation of *ϕ*_2_ condensates occurs in the dilute phase (Fig. 2a right 1). We provide a simple argument for why the kink occurs near *χ*_12_ = −1 (SM Section A5), and a recent more complex theory may provide a better estimation [24]. In this work, we primarily focus on the more interesting region where the secondary phase separation forms via interfacial instability.

**FIG. 2.**
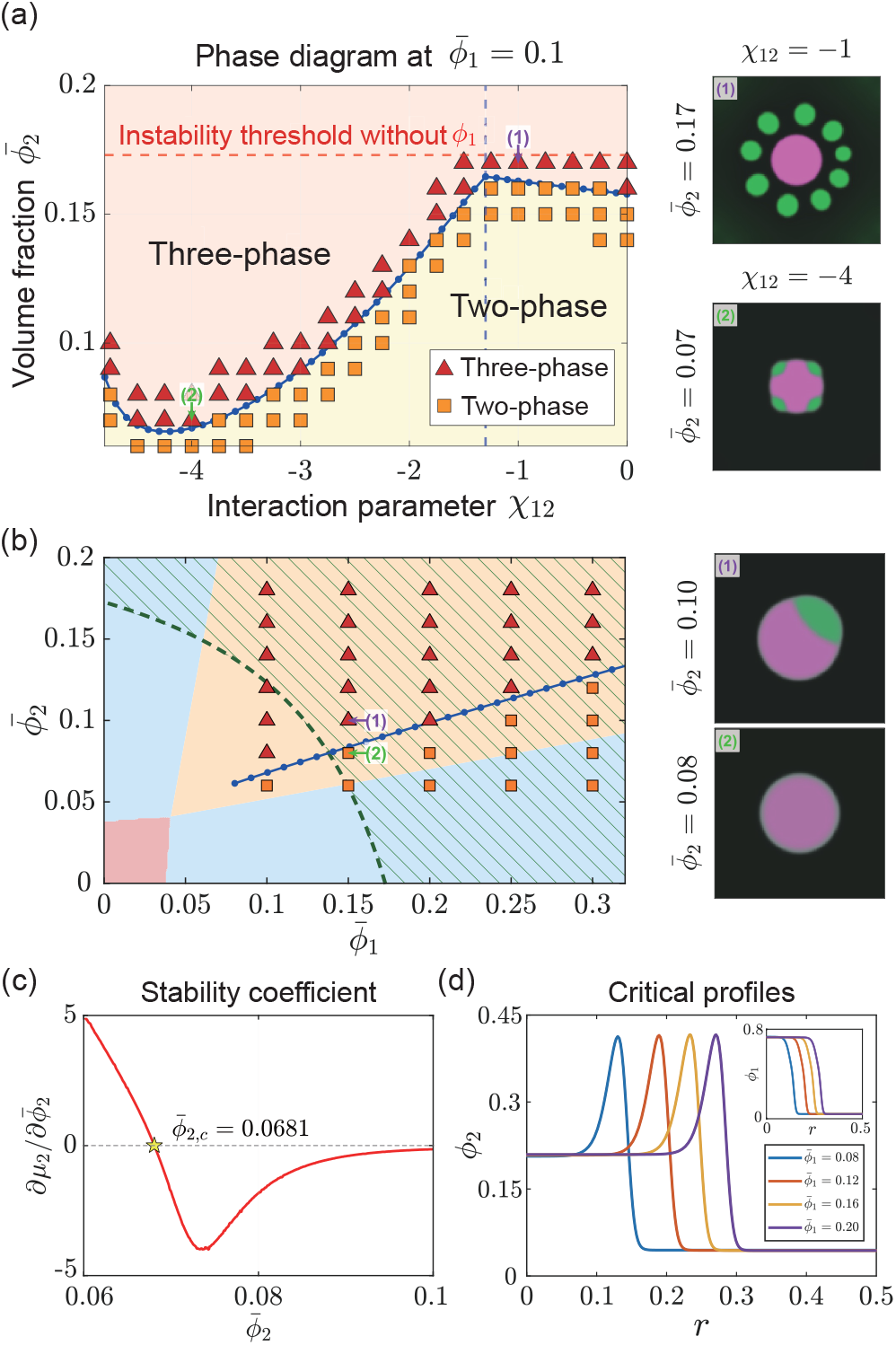
Quasistatic simulations and the predicted interfacial instability line. (a) We quasistatically increase 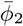 to a solution with a preexisting *ϕ*_1_ condensate and monitor when the system exhibits secondary phase separation. The markers and background colors label whether the system is in a three-phase or two-phase state. In this panel, 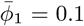. The blue line is the predicted interfacial instability line, and the red dashed line is the instability threshold in the absence of *ϕ*_1_. Notably, the preexisting condensate significantly lowers the instability threshold. Snapshots at *t* = 2*t*_*o*_ corresponding to the parameters (1) and (2) are shown on the right. Here, *t*_*o*_ marks the onset of secondary phase separation (i.e., the time at which the system exhibits three phases; see details in SM section B3). (b) Similar to panel a, but in the parameter space of 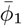 and 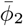. In this panel, we fix *χ*_12_ = − 4.5. The background color represents the number of phases in the corresponding ground state. Red: one phase. Blue: two phases. Yellow: three phases. The blue line is the predicted interfacial instability line. Snapshots for the parameters (1) and (2) are shown on the right, taken when the system reaches a steady state (SM section B3). (c) The stability coefficient, 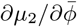_2_, and interfacial instability occur when it becomes negative. (d) The equilibrium *ϕ*_2_ profiles at the instability onset, and the inset shows the corresponding *ϕ*_1_ profiles.

We next fix *χ*_12_ and study when the transition from a two-phase state to a three-phase state occurs as 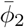 increases as a function of 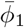 (Fig. 2b). Under the quasistatic protocol, the system remains in the metastable two-phase state even if the ground state already becomes a three-phase state until interfacial instability triggers the secondary phase transition (Fig. 2b).

To theoretically predict the onset of interfacial instability, we exploit the system’s rotational symmetry and numerically find the equilibrium profiles as a function of the radial coordinate *r* with *r* = 0 at the center of the preexisting condensate, i.e., solving Eq. (3) with the constraints 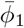 and 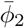 fixed at the target values (see theoretical and numerical details in SM section A4 and B2). We monitor how the chemical potential of *ϕ*_2_ changes and compute the stability coefficient 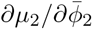 (Fig. 2c). We expect that the onset of instability should occur when 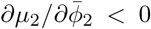. Beyond this point, the system spontaneously breaks its rotational symmetry and a new phase enriched in *ϕ*_2_ forms. Indeed, this predicted threshold 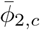 to trigger interfacial instability, which we call the interfacial instability line, precisely matches the simulations (see the blue lines in Fig. 2a and b).

Intuitively, one can imagine cutting the system into multiple sectors that can exchange mass, where each sector is locally in equilibrium. When 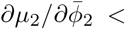0, an increase in mass on any given sector lowers its chemical potential, which drives further mass influx and leads to instability. We include a more detailed derivation for the instability criteria in SM Section A6.

Finally, we remark that as *ϕ*_1_ increases, the preexisting condensate size increases, so the amount of client protein enriched inside the condensate increases as well. Meanwhile, the *ϕ*_2_(*r*) profile when the interfacial instability occurs has a nearly invariant shape near the condensate surface (Fig. 2d). Therefore, we predict that the threshold *ϕ*_2_ to trigger instability should increase approximately linearly as a function of 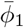, which is indeed observed in the simulations (Fig. 2b).

### Adding client rapidly

We return to the case where *ϕ*_2_ is rapidly added all at once to the solution, which can be biologically more relevant. We randomly sample 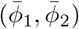 in the parameter space and count the number of phases at the end of the simulations (Fig. 3a). Strikingly, the interfacial instability line predicted by the quasistatic theory again precisely separates the two-phase and three-phase states (the blue line in Fig. 3a). The unexpected results suggest that the kinetic pathway via interfacial instability is a universal mechanism to generate three-phase coexistence in ternary mixtures (Fig. 1b and Fig. 3b2,3).

**FIG. 3.**
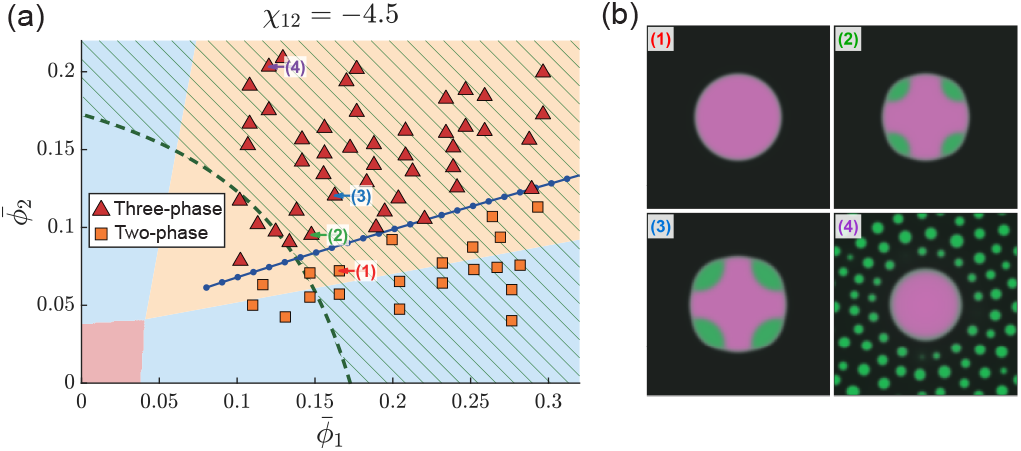
The quasistatic theory also works for the situation when *ϕ*_2_ is added rapidly. (a) Similar to the phase diagram in panel b of Fig. 2, but with *ϕ*_2_ rapidly added. We randomly sample 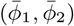 and count the number of phases at the end of the simulations. We evolve the system until it reaches a steady state or exhibits a three-phase coexistence, or up to a maximum time of *t* = 5 *×* 10^6^, whichever comes first (details of simulation termination criteria are in SM Section B3). The blue line is the predicted interfacial instability line according to the quasistatic theory. The background color represents the number of phases in the corresponding ground state. Red: one phase. Blue: two phases. Yellow: three phases. (b) Snapshots with parameters highlighted in (a). In (1), the snapshot is taken when the system reaches the steady state. In (2-4), the snapshots are at *t* = 2*t*_*o*_, where *t*_*o*_ marks the onset of secondary phase separation.

We notice that the new phase can fully enclose the preexisting condensate, bypassing the interfacial instability, if 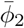 is large enough (Fig. S1). Moreover, if 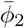 is so large that the homogeneous dilute phase is locally unstable immediately after *ϕ*_2_ addition, new condensates form directly in the dilute phase without touching the *ϕ*_1_ condensate, which was also observed experimentally (Fig. 3b4) [22].

### Rapid addition of the client and scaffold simultaneously

We next study the even more dramatic case where we rapidly add the scaffold and client protein simultaneously. At *t* = 0, the system is initialized as a homogeneous mixture. We remark that while the linear stability analysis tells us whether the initial state is stable, it is unclear what the actual steady state the system will reach.

We randomly sample 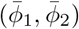 in the parameter space and count the number of phases at the end of the simulations. Surprisingly, the interfacial instability lines from the quasistatic theory and the bulk spinodal line determined from the initial state together dictate the final steady states (see the green dashed line and the blue lines in Fig. 4a). If the parameters 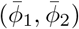 do not cross the bulk spinodal line, the uniform state will stay stable even if the ground state is a two-phase state (Fig. 4b1). Strikingly, beyond the bulk spinodal line, the interfacial instability lines again separate the two-phase and three-phase states (Fig. 4b2, 3), which is nontrivial since the theory by itself assumes a quasistatic *ϕ*_2_ addition while the simulations are clearly not. Notably, in this case, *ϕ*_1_ and *ϕ*_2_ are symmetric (Fig. 4b3, 4) so that there are two interfacial instability lines (Fig. 4a).

**FIG. 4.**
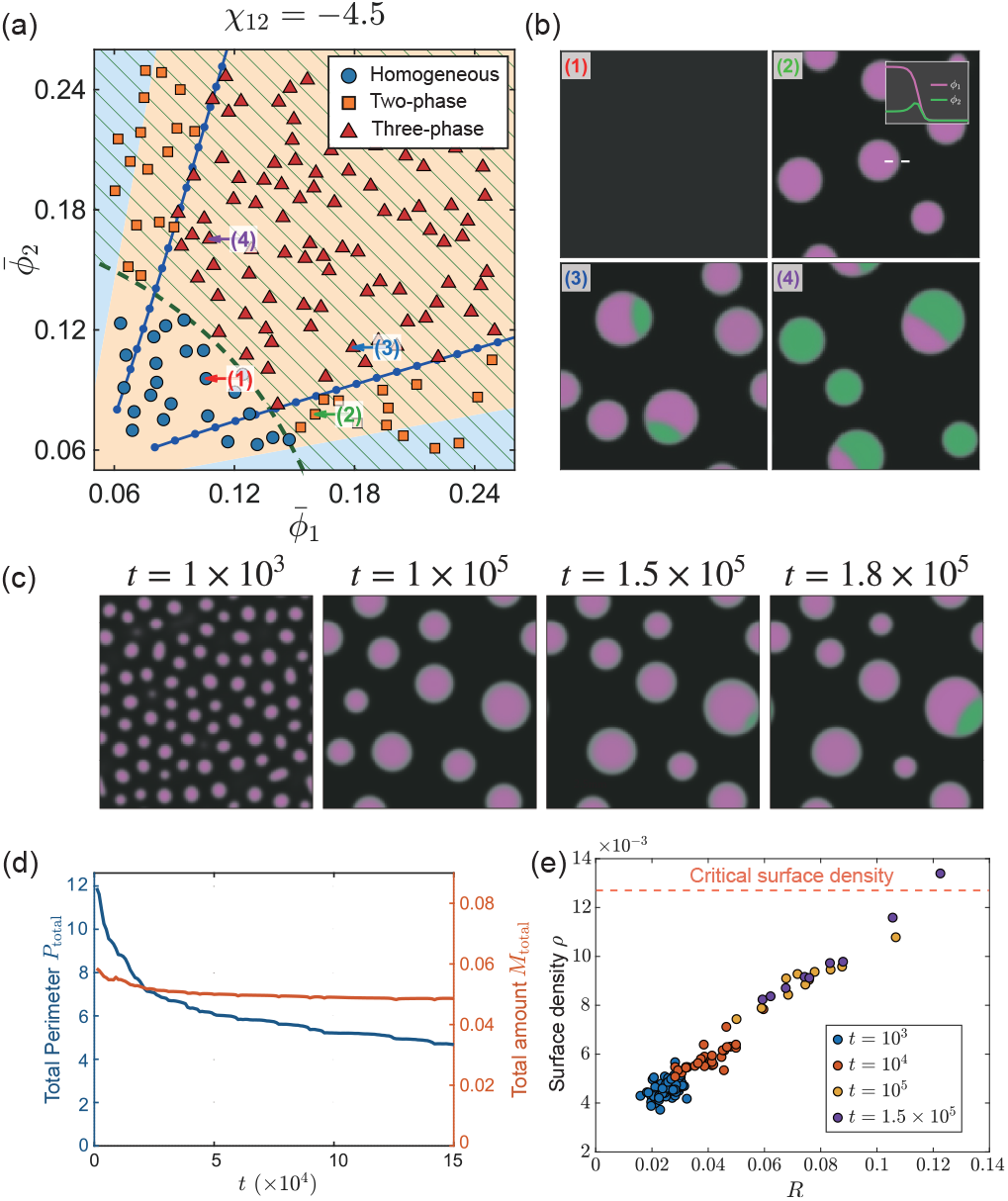
The quasistatic theory also works when the scaf-fold and client are added simultaneously. (a) Similar to the phase diagram in panel a of Fig. 3, but with both *ϕ*_1_ and *ϕ*_2_ rapidly added. We evolve the system until it reaches a steady state or exhibits three-phase coexistence, or until *t* = 5 *×* 10^6^, whichever comes first. In this case, two symmetric interfacial instability lines (the blue lines) appear. The green dashed line is the bulk spinodal line for the initial uniform state. (b) Snapshots at *t* = 2.5 *×* 10^5^ for a homogeneous state (1), a two-phase state (2), and two three-phase states (3, 4). (c) The spatiotemporal evolution of the *ϕ*_1_ and *ϕ*_2_ fields. As the coarsening process continues, interfacial instability first occurs on the largest condensate. (d) During the coarsening process, the total perimeter of all condensates decreases, while the total amount of surface-localized client protein remains approximately constant. (e) Scatter plots of the radii and the surface densities of *ϕ*_2_ for all surviving condensates at different times. The red dashed line marks the critical surface density of *ϕ*_2_ for secondary phase separation, which is approximately independent of the condensate size (Fig. S2).

The spatiotemporal evolution of the system exhibits three stages. In the first stage (the snapshot at *t* = 10^3^ in Fig. 4c), the system undergoes rapid spinodal decomposition, generating numerous primary condensates. In the second stage (compare the snapshots at *t* = 10^3^ and *t* = 10^5^), the system undergoes coarsening, during which large primary condensates grow while small ones shrink. Crucially, this coarsening process reduces the perimeters of condensates, leading to a gradual accumulation of *ϕ*_2_ around the condensate surfaces. Ultimately, in the last stage (compare the snapshots at *t* = 1.5 *×* 10^5^ and 1.8 *×* 10^5^), the accumulation of *ϕ*_2_ at the interface triggers the secondary phase separation.

To verify that interfacial instability is driven by coarsening, we compute the time evolution of the total perimeter of all condensates and the total amount of *ϕ*_2_ localized at the condensate surfaces (Fig. 4d; see numerical details in SM Section A7). Indeed, the total amount of surface-localized *ϕ*_2_ changes mildly while the total perimeter decreases significantly. We further compute the surface density of *ϕ*_2_ on each condensate (the amount of surface-localized *ϕ*_2_ divided by the perimeter). Indeed, as coarsening continues, condensates grow larger, with more *ϕ*_2_ accumulating at their surfaces (Fig. 4e).

Notably, the surface density of *ϕ*_2_ correlates positively with the condensate size (Fig. 4e). This size dependence can be attributed to the lower interfacial curvature of larger condensates, which lowers the local chemical potential of *ϕ*_2_ (see SM Section A8 for a detailed derivation). Consequently, the largest condensate, possessing the highest *ϕ*_2_ surface density, is the first to cross the instability threshold and trigger secondary phase separation, consistent with our quasistatic theory (Fig. 2d; see snapshots at *t* = 1.5 *×* 10^5^ and 1.8 *×* 10^5^ in Fig. 4c).

## Discussion

In this work, we unveil a universal kinetic pathway to generate a new phase in a ternary mixture: interfacial instability at the surfaces of preexisting condensates. This pathway requires significantly less client protein to form the new phase than in a system without preexisting condensates. By studying the quasistatic protocol and using the system’s rotational symmetry, we successfully predict when interfacial instability occurs, characterized by a negative thermodynamic stability coefficient.

Strikingly, the quasistatic theory even predicts whether secondary phase separation can occur when the client is added rapidly, or when both the client and the scaffold are added rapidly. The robustness of the quasistatic theory suggests that interfacial instability is a universal pathway for generating a new phase. Notably, when both the client and scaffold are added rapidly, the coarsening process continuously decreases the perimeters of all condensates so that the surface densities of the client gradually increase, eventually triggering the interfacial instability. Meanwhile, because the surface density correlates positively with condensate size, the new phase always nucleates first at the largest condensate.

Our work provides a physical explanation for the formation of TDP-43 puncta on the surface of stress granules both *in vivo* and *in vitro* [22], suggesting that our conclusions generalize to more complex solutions with more than three components. Therefore, we expect that the interface-mediated pathway for generating new phases should be generally applicable beyond ternary mixtures. Indeed, similar adsorption-to-demixing transitions have been observed in synthetic polymer systems with four components [25]. Our work also implies that living cells can actively exploit preexisting condensates to promote the spatially targeted phase separation of essential proteins. By doing so, cells can effortlessly bypass high nucleation barriers without incurring the energetic cost of protein upregulation.

Meanwhile, such surface-localized client-rich phases may create highly concentrated environments that favor subsequent irreversible, sometimes disease-associated aggregation [2, 3, 22, 26–29]. We speculate that surface clusters on some biomolecular condensates, e.g., MEG3 clusters on P-granule surfaces, may also form through this interface-mediated mechanism [30, 31]. By tuning the interaction between the client and the scaffold via chaperones or small molecules, it may be possible to arrest pathological aggregation at the benign surface-enrichment stage.

We thank Sheng Mao and Zhiheng Wang for helpful discussion related to this work. The research was funded by National Natural Science Foundation of China (Grant No. 12474190), National Key Research and Development Program of China (2024YFA0919600), and Peking-Tsinghua Center for Life Sciences grants.

