## Supplementary Material for "Interface-mediated secondary phase separation"

### Supplemental Material

Kaifeng Weng

*Center for Quantitative Biology, Peking University, Beijing, China and*
*School of Physics, Peking University, Beijing, China*

Jie Lin

*Center for Quantitative Biology, Peking University, Beijing, China and*
*Peking-Tsinghua Center for Life Sciences, Peking University, Beijing, China*

(Dated: September 4, 2026)

#### A. THEORETICAL MODEL

##### A1. Dynamic Framework: The Multicomponent Cahn-Hilliard Equations

To rigorously derive the dynamic evolution equations for our system, we employ the variational principle based on the Rayleigh dissipation formalism [1]. Consider a general system with $N+1$ components. We denote the local volume fraction of the $i$-th component as $\phi_i$, which is subject to the incompressibility constraint $\sum_{i=1}^{N+1} \phi_i = 1$. The total free energy of the system is given by

$$F^* = \int f^*(\phi_i, \nabla\phi_i) dV, \quad i = 1, 2, \cdots, N+1. \tag{1}$$

where $f^*(\phi_i, \nabla\phi_i)$ represents the free energy density. Throughout this text, the asterisk ($*$) designates thermodynamic quantities defined in the explicit-solvent frame retaining all $N+1$ compositional degrees of freedom, i.e., the volume fraction of the solvent $\phi_{N+1}$ is treated as an independent variable.

Given the equivalence between the solutes and the solvent, the Rayleighian of the system can be constructed as follows:

$$R = \int \sum_{i=1}^{N+1} \left[ \frac{\delta F^*}{\delta \phi_i} \dot{\phi}_i + \frac{1}{4} \zeta \sum_{j=1}^{N+1} \phi_i \phi_j (v_i - v_j)^2 + p\phi_i v_i \right] dV, \tag{2}$$

where $v_i$ is the velocity of component $i$, $\zeta$ is the friction coefficient density between different components, which is assumed to be a constant. We note that the time evolution $\dot{\phi}_i$ satisfies the continuity equation $\dot{\phi}_i = -\nabla \cdot (\phi_i v_i)$. Therefore, the incompressibility constraint $\sum_{i=1}^{N+1} \phi_i = 1$, combined with the assumption of no macroscopic flow, implies that $\sum_j \phi_j v_j = 0$. To consider this effect, we introduce $p$ as a Lagrange multiplier to enforce the constraint.

Replacing $\dot{\phi}_i$ by $-\nabla \cdot (\phi_i v_i)$ in Eq. (2) and minimizing the Rayleighian with respect to the velocity field ($\delta R / \delta v_i = 0$), we obtain:

$$\phi_i \nabla \left( \frac{\delta F^*}{\delta \phi_i} \right) + \zeta \phi_i v_i + p\phi_i = 0. \tag{3}$$

Summing over $i$ from 1 to $N+1$, the Lagrange multiplier can be explicitly determined as

$$p = -\sum_i \phi_i \nabla \left( \frac{\delta F^*}{\delta \phi_i} \right). \tag{4}$$

Substituting $p$ back yields the velocity field:

$$v_i = -\frac{1}{\zeta}\left[\nabla\frac{\delta F^*}{\delta\phi_i} - \sum_j \phi_j \nabla\frac{\delta F^*}{\delta\phi_j}\right]. \tag{5}$$

Inserting $v_i$ into the continuity equation $\frac{\partial\phi_i}{\partial t} = -\nabla\cdot(\phi_i v_i)$, we arrive at the dynamic equation:

$$\frac{\partial\phi_i}{\partial t} = \nabla\cdot\left[\sum_{j=1}^{N+1}\Lambda_{ij}\nabla\frac{\delta F^*}{\delta\phi_j}\right], \text{ where } \Lambda_{ij} = \frac{1}{\zeta}\phi_i(\delta_{ij}-\phi_j). \tag{6}$$

In this work, we further simplify the model by treating the $(N+1)$-th component (solvent) as $\phi_{N+1} = 1 - \sum_{i=1}^{N}\phi_i$ and redefine the free energy functional as a function of $\phi_i$ for $i = 1, 2, ..., N$:

$$F(\phi_1, \phi_2, \cdots, \phi_N) = F^*(\phi_1, \cdots, \phi_N, 1 - \sum_{i=1}^{N}\phi_i). \tag{7}$$

Therefore, the functional derivative of $F$ becomes:

$$\frac{\delta F}{\delta\phi_i} = \frac{\delta F^*}{\delta\phi_i} - \frac{\delta F^*}{\delta\phi_{N+1}}. \tag{8}$$

By substituting this into Eq. (6), we find the Cahn-Hilliard equation governing the spatial-temporal evolution of the $N$ independent variables as

$$\frac{\partial\phi_i}{\partial t} = \nabla\cdot\left[\sum_{j=1}^{N}\Lambda_{ij}\nabla\frac{\delta F}{\delta\phi_j}\right] \quad (i = 1, \ldots, N). \tag{9}$$

Interesting, the $\Lambda$ matrix remains the same form $\Lambda_{ij} = \phi_i(\delta_{ij} - \phi_j)/\zeta$ while the index now runs from 1 to $N$. For ternary mixtures, $N = 2$. We next introduce the explicit form of the free energy functional.

##### A2. Thermodynamic Model: Flory-Huggins Free Energy for Ternary Mixtures

To describe this multi-component system, we employ the classical Flory-Huggins model with a gradient penalty term. For an $(N + 1)$-component mixture, the free energy density $f^*$ is given by:

$$f^* = \frac{k_B T}{v_0}\left[\sum_{i=1}^{N+1}\phi_i\ln\phi_i + \frac{1}{2}\sum_{i,j=1}^{N+1}\chi^*_{ij}\phi_i\phi_j - \frac{\lambda^2}{2}\sum_{i,j=1}^{N+1}\chi^*_{ij}\nabla\phi_i\cdot\nabla\phi_j\right], \tag{10}$$

where $\chi^*_{ij}$ is a symmetric matrix with $\chi^*_{ii} = 0$.

Using the incompressibility constraint $\sum_{i=1}^{N+1} \phi_i = 1$, we can map the free energy density to a new one with $N$ independent variables, i.e., $\phi_i$ where $i$ runs from 1 to $N$. For example,

$$\frac{1}{2}\sum_{i=1}^{N+1}\sum_{j=1}^{N+1} \chi^*_{ij}\phi_i\phi_j \to \frac{1}{2}\sum_{i=1}^{N}\sum_{j=1}^{N}(\chi^*_{ij} - \chi^*_{i,N+1} - \chi^*_{N+1,j})\phi_i\phi_j. \tag{11}$$

In the above equation, we neglect linear terms as they merely shift the chemical potentials by a constant and do not alter thermodynamic equilibrium. Therefore, we define the effective interaction matrix as

$$\chi_{ij} = \chi^*_{ij} - \chi^*_{i,N+1} - \chi^*_{N+1,j}. \tag{12}$$

The free energy density functional $f(\phi_1, \ldots, \phi_N)$, excluding the explicit solvent variable, becomes

$$f = \frac{k_B T}{v_0}\left[\sum_{i=1}^{N}\phi_i \ln\phi_i + \left(1 - \sum_{i=1}^{N}\phi_i\right)\ln\left(1 - \sum_{i=1}^{N}\phi_i\right) + \frac{1}{2}\sum_{i,j=1}^{N}\chi_{ij}\phi_i\phi_j - \frac{\lambda^2}{2}\sum_{i,j=1}^{N}\chi_{ij}\nabla\phi_i\cdot\nabla\phi_j\right]. \tag{13}$$

The functional derivative of the system's total free energy becomes

$$\frac{\delta F}{\delta\phi_i} = \frac{k_B T}{v_0}\left[\ln\left(\frac{\phi_i}{1 - \sum_k^N \phi_k}\right) + \sum_{j=1}^{N}\chi_{ij}\phi_j + \lambda^2\sum_{j=1}^{N}\chi_{ij}\nabla^2\phi_j\right] = \frac{k_B T}{v_0}\tilde{\mu}_i, \tag{14}$$

where $\tilde{\mu}$ is the dimensionless exchange chemical potential. Substituting this explicit form of $\delta F/\delta\phi_i$ into the multi-component Cahn-Hilliard equations established in Section A1, we obtain

$$\frac{\partial\phi_i}{\partial t} = \nabla\cdot\left[\sum_{j=1}^{N}\frac{k_B T}{v_0}\Lambda_{ij}\nabla\tilde{\mu}_j\right] \quad (i = 1, \ldots, N). \tag{15}$$

To characterize the system's kinetic transport, we define the mobility matrix as:

$$M_{ij} = \frac{k_B T}{v_0}\Lambda_{ij} = D\phi_i(\delta_{ij} - \phi_j), \tag{16}$$

where $D = k_B T/(v_0\zeta)$ is the diffusion constant. To facilitate numerical simulations, we cast the dynamical equations into a dimensionless form in the following section.

##### A3. Nondimensionalization

To nondimensionalize the system, we use the system size $L_0$ as the length unit so that the dimensionless coordinate $\tilde{x} = x/L_0$, and the corresponding gradient operator becomes

$\tilde{\nabla} = L_0 \nabla$. The expression for the dimensionless exchange chemical potential becomes:

$$\tilde{\mu}_i = \ln\left(\frac{\phi_i}{1-\phi_1-\phi_2}\right) + \sum_{j=1}^{2} \chi_{ij}\phi_j + \tilde{\lambda}^2 \sum_{j=1}^{2} \chi_{ij} \tilde{\nabla}^2 \phi_j, \tag{17}$$

where $\tilde{\lambda} = \lambda / L_0$ defines the dimensionless interfacial width.

For the temporal evolution, we set the time unit as $\tau = \lambda^2/D$, which physically represents the time required for a molecule to diffuse across the interfacial width $\lambda$. Scaling the time variable as $\tilde{t} = t/\tau$ yields the fully nondimensionalized dynamical equations:

$$\frac{\partial \phi_i}{\partial \tilde{t}} = \tilde{\lambda}^2 \tilde{\nabla} \cdot \left( \sum_{j=1}^{2} \tilde{M}_{ij} \tilde{\nabla} \tilde{\mu}_j \right), \tag{18}$$

where $\tilde{M}_{ij} = \phi_i(\delta_{ij} - \phi_j)$ is the dimensionless mobility matrix, and the indices denote the two independent solute components ($i = 1, 2$). We also set the unit of the free energy density to $k_B T / v_0$. For notational simplicity, we remove the tildes on the dimensionless variables in the main text and in the following Supplemental Material. All variables are dimensionless unless otherwise noted.

##### A4. Reduction to the One-Dimensional Model

In this section, we discuss the model for a rotationally symmetric system, so all variables depend solely on the radial coordinate $r$ in polar coordinates. For simplicity, we model the system geometry as a circle with radius $L$, and the total free energy of the system becomes:

$$F[\phi_1, \phi_2] = 2\pi \int_0^L \left[ f_b(\phi_1, \phi_2) - \frac{1}{2}\lambda^2 \sum_{i,j=1}^{2} \chi_{ij} \nabla\phi_i \cdot \nabla\phi_j \right] r dr,$$

where the spatial gradient simplifies to $\nabla = \hat{e}_r \frac{d}{dr}$, and $f_b$ denotes the homogeneous Flory-Huggins free energy density:

$$f_b(\phi_1, \phi_2) = \phi_1 \ln \phi_1 + \phi_2 \ln \phi_2 + (1-\phi_1-\phi_2)\ln(1-\phi_1-\phi_2) + \frac{1}{2}\sum_{i,j=1}^{2} \chi_{ij}\phi_i\phi_j. \tag{19}$$

To account for total mass conservation of each component, we introduce the Lagrange multipliers $\mu_i$ ($i = 1, 2$) and construct the constrained functional:

$$G = F - \sum_{i=1}^{2} \mu_i \left( 2\pi \int_0^L \phi_i \, r dr - A\bar{\phi}_i \right), \tag{20}$$

where $A = \pi L^2$ is the total area of the circular domain and $\bar{\phi}_i$ is the mean volume fraction. At thermodynamic equilibrium, $\delta G/\delta\phi_i = 0$, which yields:

$$\frac{\delta G}{\delta \phi_i} = \frac{\delta F}{\delta \phi_i} - \mu_i = 0. \tag{21}$$

Thus, we have

$$\frac{\delta F}{\delta \phi_i} = \mu_{i,b}(r) + \lambda^2 \sum_{j=1}^{2} \chi_{ij} \left[ \phi_j''(r) + \frac{1}{r}\phi_j'(r) \right] = \mu_i, \quad (i = 1, 2), \tag{22}$$

where $\mu_{i,b}(r) \equiv \partial f_b/\partial \phi_i = \ln \phi_i - \ln(1 - \phi_1 - \phi_2) + \sum_{j=1}^{2} \chi_{ij}\phi_j$ represents the local bulk chemical potential. Solving the equilibrium conditions explicitly by inverting the interaction matrix, we obtain:

$$\begin{aligned} \phi_1'' + \frac{1}{r}\phi_1' &= \frac{\chi_{22}\left[\mu_1 - \mu_{1,b}(r)\right] - \chi_{12}\left[\mu_2 - \mu_{2,b}(r)\right]}{\lambda^2(\chi_{11}\chi_{22} - \chi_{12}^2)} \\ \phi_2'' + \frac{1}{r}\phi_2' &= \frac{\chi_{11}\left[\mu_2 - \mu_{2,b}(r)\right] - \chi_{12}\left[\mu_1 - \mu_{1,b}(r)\right]}{\lambda^2(\chi_{11}\chi_{22} - \chi_{12}^2)}, \end{aligned} \tag{23}$$

which are subject to zero-flux boundary conditions at both the origin ($r = 0$) and the boundary ($r = L$):

$$\left.\frac{d\phi_i}{dr}\right|_{r=0} = 0, \quad \left.\frac{d\phi_i}{dr}\right|_{r=L} = 0, \quad (i = 1, 2). \tag{24}$$

The Lagrange multiplier $\mu_i$ is determined by the total mass conservation:

$$A\bar{\phi}_i = 2\pi \int_0^L \phi_i(r) r \, dr \quad (i = 1, 2). \tag{25}$$

This formulation casts the system's equilibrium structure into a standard boundary value problem (BVP), providing the theoretical foundation for the numerical integration described in section B2.

##### A5. An Argument on the Threshold $\chi_{12}$ Above Which $\phi_2$ is Excluded From the Preexisting Condensate

In the main text, we show that for $\chi_{12} \gtrsim -1$, the preexisting condensate simply acts as an excluded volume. To understand the physical origin of the threshold $\chi_{12}$ value, we neglect the gradient energy and consider the bulk chemical potential of $\phi_2$:

$$\mu_2 = \ln(\phi_2) - \ln(1 - \phi_1 - \phi_2) + \chi_{12}\phi_1 + \chi_{22}\phi_2. \tag{26}$$

We argue that for $\phi_2$ to move from the dilute phase to the dense phase spontaneously, the chemical potential of $\phi_2$ must decrease as the local $\phi_1$ concentration increases. This requires $\partial\mu_2/\partial\phi_1|_{\phi_1=0} < 0$, which yields the following inequality:

$$\left.\frac{\partial\mu_2}{\partial\phi_1}\right|_{\phi_1=0} = \frac{1}{1-\phi_2} + \chi_{12} < 0. \tag{27}$$

This inequality defines the critical interaction parameter threshold for adsorption:

$$\chi_{12}^{\text{critical}} = -\frac{1}{1-\phi_2}. \tag{28}$$

The term $\frac{1}{1-\phi_2}$ represents the entropic penalty, while $\chi_{12}$ captures the enthalpic attraction. Adsorption occurs only when the attraction overcomes this entropic penalty ($\chi_{12} < \chi_{12}^{\text{critical}}$). Notably, for $\phi_2 \ll 1$, this threshold naturally simplifies to:

$$\chi_{12}^{\text{critical}} \approx -1. \tag{29}$$

This explains why the kink in Fig. 2a of the main text occurs around $\chi_{12} = -1$.

##### A6. Detailed derivation for the Instability Threshold

We define the integrated mass of the $\phi_2$ field along a radial profile at a given azimuthal angle $\theta$ as:

$$m_2(\theta) = \int_0^L \phi_2(r,\theta) r\, dr. \tag{30}$$

We adopt a local equilibrium approximation and assume that for a given mass $m_2(\theta)$, the radial profile instantaneously relaxes to a steady state, denoted by:

$$\phi_i(r,\theta) = \phi_i^s\big(r, m_2(\theta)\big). \tag{31}$$

The total free energy of the system can be written in polar coordinates as:

$$F = \int_0^{2\pi}\int_0^L \left[ f_b(\phi_1^s, \phi_2^s) + \frac{\lambda^2}{2}\sum_{i,j}\chi_{ij}\partial_r\phi_i^s\partial_r\phi_j^s + \frac{\lambda^2}{2r^2}\sum_{i,j}\chi_{ij}\partial_\theta\phi_i^s\partial_\theta\phi_j^s \right] r\, dr\, d\theta. \tag{32}$$

The first two terms constitute the total radial free energy, $F_r$, which depends solely on the angular mass distribution $m_2(\theta)$:

$$F_r = \int_0^{2\pi}\int_0^L \left[ f_b + \frac{\lambda^2}{2}\sum_{i,j}\chi_{ij}\partial_r\phi_i^s\partial_r\phi_j^s \right] r\, dr\, d\theta = \int_0^{2\pi} g\left[m_2(\theta)\right] d\theta, \tag{33}$$

where $g[m_2]$ represents the radial free energy density per unit angle. The final term is the azimuthal gradient free energy, $F_a$. By applying the chain rule $\partial_\theta \phi_i^s = (\partial \phi_i^s / \partial m_2) m_2'(\theta)$, this term becomes:

$$F_a = \int_0^{2\pi} \int_0^L \left[ \frac{\lambda^2}{2r^2} \sum_{i,j} \chi_{ij} \partial_\theta \phi_i^s \partial_\theta \phi_j^s \right] r \, dr \, d\theta = \int_0^{2\pi} \frac{1}{2} \Gamma \left( m_2(\theta) \right) \left( m_2'(\theta) \right)^2 d\theta. \tag{34}$$

Here, we define $\psi_i(r, m_2) \equiv \partial \phi_i^s(r, m_2) / \partial m_2$, and introduce the effective angular gradient coefficient:

$$\Gamma \left( m_2(\theta) \right) = \int_0^L \left[ \frac{\lambda^2}{r} \sum_{i,j} \chi_{ij} \psi_i(r, m_2) \psi_j(r, m_2) \right] dr. \tag{35}$$

The total free energy of the system is then compactly written as a functional of $m_2(\theta)$:

$$F_{total}[m_2] = \int_0^{2\pi} \left[ g\big(m_2(\theta)\big) + \frac{1}{2} \Gamma\big(m_2(\theta)\big) \left( m_2'(\theta) \right)^2 \right] d\theta. \tag{36}$$

We now consider a base state with an axisymmetric mass distribution $m_0$ and introduce a small perturbation $\delta m(\theta)$, yielding:

$$m_2(\theta) = m_0 + \delta m(\theta). \tag{37}$$

Expanding the free energy to the second order in $\delta m$, and noting that the first-order term vanishes due to mass conservation ($\int_0^{2\pi} \delta m(\theta) d\theta = 0$), the variation in the free energy is:

$$\Delta F = \frac{1}{2} \int_0^{2\pi} \left[ g''(m_0) \delta m(\theta)^2 + \Gamma(m_0) \left( \delta m'(\theta) \right)^2 \right] d\theta. \tag{38}$$

By decomposing the angular mass perturbation into Fourier modes, $\delta m(\theta) = A \cos(n\theta)$ with an integer wavenumber $n \geq 1$, the integral evaluates to:

$$\Delta F = \frac{\pi A^2}{2} \Big[ g''(m_0) + n^2 \Gamma(m_0) \Big]. \tag{39}$$

The criterion for instability is $\Delta F < 0$.

Note that $g''(m_0)$ represents the derivative of the chemical potential with respect to mass, $g''(m_0) = \partial \mu_2 / \partial m_2$; the condition becomes:

$$\frac{\pi A^2}{2} \left[ \frac{\partial \mu_2}{\partial m_2} + n^2 \Gamma(m_0) \right] < 0. \tag{40}$$

Denoting the averaged volume fraction as $\bar{\phi}_2$, we can rewrite the instability criterion as:

$$\frac{2}{L^2} \frac{\partial \mu_2}{\partial \bar{\phi}_2} + n^2 \Gamma(m_0) < 0. \tag{41}$$

Since $m_2$ scales as $L^2$, the gradient penalty term $\Gamma(m_0)$ scales as $\lambda^2/L^4$, which is usually much smaller than the term $\frac{2}{L^2}\frac{\partial\mu_2}{\partial\bar{\phi}_2}$. We can therefore neglect it, obtaining the simplified instability criterion:

$$\frac{\partial\mu_2}{\partial\bar{\phi}_2} < 0. \tag{42}$$

However, we note that when $\bar{\phi}_2$ is high enough, this gradient penalty term can become nonnegligible and suppress the instability (Fig. S1).

##### A7. Quantification of the Amount of $\phi_2$ Localized at Condensate Surfaces

To quantitatively track the accumulation of the client component ($\phi_2$) on the surfaces of the primary condensates, we construct an interfacial weight function $\mathcal{W}(\phi_1)$ that effectively isolates the interfaces from other regions:

$$\mathcal{W}(\phi_1) = \frac{4(\phi_1 - \phi_1^{\min})(\phi_1^{\max} - \phi_1)}{(\phi_1^{\max} - \phi_1^{\min})^2}, \tag{43}$$

where $\phi_1^{\min}$ and $\phi_1^{\max}$ correspond to the equilibrium bulk volume fractions of the dilute and dense phases, respectively, which remain approximately constant across different condensates. This weight function naturally peaks at unity at the center of the interface ($\phi_1 = \frac{\phi_1^{\max}+\phi_1^{\min}}{2}$) and decays to zero in both bulk phases.

Using this weight function, the total accumulated amount of $\phi_2$ at the interfaces across the entire domain $\Omega$ can be evaluated via the integral:

$$N_2^{\text{tot}} = \int_\Omega \mathcal{W}(\phi_1)\phi_2\, dA. \tag{44}$$

Meanwhile, we also compute the perimeter for each condensate. Specifically, we compute the area for each condensate where sites with $\phi > (\phi_1^{\max} + \phi_1^{\min})/2$ are considered as part of the condensate; from the area, we infer the perimeter. We then compute the total perimeter of all condensates, $L^{\text{tot}}$.

As demonstrated in Fig. 4d of the main text, during the late-stage coarsening process, the total perimeter $L^{\text{tot}}$ monotonically decreases. In contrast, the total accumulated amount of $\phi_2$ at the interfaces, $N_2^{\text{tot}}$, remains nearly constant, which inevitably leads to an increasingly higher surface density of $\phi_2$ along the interfaces.

For each individual condensate $k$ with perimeter $L^{(k)}$, we also compute the local surface density as:

$$\rho_2^{(k)} = \frac{1}{L^{(k)}} \int_{\Omega_k} \mathcal{W}(\phi_1)\phi_2 \, dA, \tag{45}$$

where $\Omega_k$ represents the local subdomain surrounding the $k$-th condensate. As the system coarsens, the surface densities, $\rho_2^{(k)}$, of individual condensates increase continuously. Furthermore, as corroborated by our numerical observations (see Fig. 4e of the main text), the local surface density $\rho_2^{(k)}$ exhibits a clear positive correlation with the condensate radius. This size-dependent accumulation is fundamentally driven by a curvature-induced chemical potential gradient, a thermodynamic mechanism we detail in the next section.

As observed in Figure 2d of the main text, the critical volume profiles of $\phi_2$ at the instability threshold exhibit nearly identical shapes for different values of $\bar{\phi}_1$. Consequently, these profiles share consistent surface densities, a phenomenon further corroborated in Figure S2. Ultimately, when the local surface density $\rho_2^{(k)}$ of a condensate exceeds this critical threshold, the secondary phase separation occurs, which always happens first for the largest condensate.

##### A8. Thermodynamic Origin of the Size-Dependent Interfacial Accumulation of $\phi_2$

To understand why larger condensates have higher surface densities of $\phi_2$ in the simulations where we rapidly add the scaffold and client simultaneously, we analyze the chemical potential of the client component:

$$\mu_2 = \ln\left(\frac{\phi_2}{1-\phi_1-\phi_2}\right) + \sum_{j=1}^{2} \chi_{2j}\phi_j + \lambda^2 \sum_{j=1}^{2} \chi_{2j} \nabla^2 \phi_j. \tag{46}$$

Given an axisymmetric spherical condensate, the volume-fraction fields are functions of the radial coordinate $r$. Recall that in polar coordinates, the Laplacian operator is:

$$\nabla^2 \phi_j = \frac{d^2\phi_j}{dr^2} + \frac{1}{r}\frac{d\phi_j}{dr}. \tag{47}$$

Given that the shape of this profile remains approximately identical across condensates with different sizes at the instability threshold (as illustrated in Fig. 2d of the main text), the chemical potential at the interface of a condensate ($r = R_k$) can be decomposed into a constant term and a curvature-dependent contribution:

$$\mu_2(R_k) \approx \mu_2^{\text{flat}} + \frac{\lambda^2}{R_k} \sum_{j=1}^{2} \chi_{2j}\phi_j'(R_k), \tag{48}$$

where $\mu_2^{\text{flat}}$ contains the bulk terms and the second-derivative term ($\phi_j''$), which are independent of condensate size. Therefore, the term $\left[\lambda^2 \sum_{j=1}^{2} \chi_{2j}\phi_j'(R_k)\right] \Big/ R_k$ acts as a curvature penalty.

Because $\phi_2$ accumulates at the interface, its local gradient vanishes at the center of the interfacial layer ($\phi_2'(R_k) \approx 0$). In contrast, $\phi_1$ decreases monotonically at the interface, exhibiting a steep negative gradient. Consequently, the penalty due to interface curvature is dominated by the term $\lambda^2\chi_{21}\phi_1'/R_k$, which is positive since $\chi_{21} < 0$. Consequently, the chemical potential $\mu_2$ at the interface of a small condensate is higher than that of a large condensate. Therefore, the client molecule will spontaneously diffuse from small condensates to large condensates. This curvature-driven diffusion produces the positive correlation between condensate radius and local surface density $\rho_2^{(k)}$ observed in our simulations.

#### B. NUMERICAL METHOD

##### B1. Semi-Implicit Fourier Spectral Method

All of our 2D computational domains employ periodic boundary conditions in both spatial directions. To efficiently solve the dimensionless multi-component Cahn-Hilliard equations derived in Section A, we employ a semi-implicit Fourier spectral method based on implicit-explicit (IMEX) time integration schemes [2, 3]. A standard explicit time integration scheme is severely restricted by the numerical stiffness arising from the fourth-order spatial derivatives. Transforming the system into Fourier space allows spatial derivatives to be evaluated as simple algebraic multiplications, significantly improving computational accuracy.

By introducing the Fourier transform $\mathcal{F}$, the volume fraction field $\phi_i(\mathbf{r}, t)$ is mapped to $\phi_{i,\mathbf{k}}(t)$, where spatial derivatives transform as $\nabla \to i\mathbf{k}$ and $\nabla^2 \to -k^2$. The chemical potential in Fourier space is formulated as:

$$\mu_{i,\mathbf{k}} = \mathcal{F}\left\{\ln\left(\frac{\phi_i}{1-\phi_1-\phi_2}\right) + \chi_{ij}\phi_j\right\}_{\mathbf{k}} + \lambda^2\chi_{ij}k^2\phi_{j,\mathbf{k}}. \tag{49}$$

The real-space chemical potential gradient is first evaluated via the inverse transform, $\nabla\mu_j = \mathcal{F}^{-1}\{i\mathbf{k}\mu_{j,\mathbf{k}}\}_{\mathbf{k}}$. The spatial flux is then formed in real space as $J_i = M_{ij}\nabla\mu_j$, whose divergence in Fourier space simplifies to:

$$\mathcal{F}\{\nabla \cdot J_i\}_{\mathbf{k}} = i\mathbf{k} \cdot \mathcal{F}\{J_i\}_{\mathbf{k}}. \tag{50}$$

To suppress numerical instabilities driven by nonlinear terms, we introduce a linear stabilizing term $A\lambda^4\nabla^4\phi_i$ [3]. Treating this term explicitly at time step $n$ and implicitly at step $n+1$, the semi-implicit time-discretized equation reads:

$$\frac{\phi_{i,\mathbf{k}}^{n+1} - \phi_{i,\mathbf{k}}^{n}}{\Delta t} = \lambda^2 i\mathbf{k} \cdot \mathcal{F}\{J_i^n\}_{\mathbf{k}} + A\lambda^4 k^4 \phi_{i,\mathbf{k}}^{n} - A\lambda^4 k^4 \phi_{i,\mathbf{k}}^{n+1}, \tag{51}$$

where the stabilization constant $A = \max(|\chi_{ij}|)/2$ is chosen, following the numerical approach adapted for multi-component systems [4], to ensure that the implicit term is sufficiently dominant to suppress numerical instabilities arising from the nonlinear dynamics.

Rearranging terms yields the update scheme:

$$\phi_{i,\mathbf{k}}^{n+1} = \frac{\phi_{i,\mathbf{k}}^{n} + \Delta t \left[\lambda^2 i\mathbf{k} \cdot \mathcal{F}\{J_i^n\}_{\mathbf{k}} + A\lambda^4 k^4 \phi_{i,\mathbf{k}}^{n}\right]}{1 + A\lambda^4 k^4 \Delta t}. \tag{52}$$

Details of the 2D simulations are included in Table S2.

##### B2. Solving the Mass-Conserved Boundary Value Problem (BVP)

As derived in Section A4, the 1D equilibrium profiles are governed by a system of second-order ODEs:

$$\begin{aligned} \phi_1'' + \frac{1}{r}\phi_1' &= \frac{\chi_{22}\left[\mu_1 - \mu_{1,b}(r)\right] - \chi_{12}\left[\mu_2 - \mu_{2,b}(r)\right]}{\lambda^2(\chi_{11}\chi_{22} - \chi_{12}^2)}, \\ \phi_2'' + \frac{1}{r}\phi_2' &= \frac{\chi_{11}\left[\mu_2 - \mu_{2,b}(r)\right] - \chi_{12}\left[\mu_1 - \mu_{1,b}(r)\right]}{\lambda^2(\chi_{11}\chi_{22} - \chi_{12}^2)}. \end{aligned} \tag{53}$$

Determining the equilibrium profile requires simultaneously solving for $\phi_i(r)$ and the unknown chemical potentials $\mu_i$, subject to zero-flux Neumann boundary conditions and total mass conservation $M_i = 2\pi \int_0^L \phi_i(r) r dr$.

To handle the integral constraints efficiently, we recast the integral mass constraints into a differential form. We introduce an 8-dimensional vector $\mathbf{Y}(r) = (y_1, \ldots, y_8)^T$, defined as:

$$\begin{gathered} y_1 = \phi_1, \quad y_2 = \phi_1', \quad y_3 = \phi_2, \quad y_4 = \phi_2', \\ y_5 = \int_0^r \phi_1(s) s ds, \quad y_6 = \int_0^r \phi_2(s) s ds, \quad y_7 = \mu_1, \quad y_8 = \mu_2. \end{gathered} \tag{54}$$

Here, the variables $y_5$ and $y_6$ represent the cumulative mass integrals, while $y_7$ and $y_8$ enforce spatial uniformity of the equilibrium chemical potentials ($\mu_i' = 0$). The original system is thus reformulated into a closed system of 8 first-order ODEs:

$$\mathbf{Y}' = \begin{pmatrix} y_2 \\ -\frac{1}{r}y_2 + f_1(y_1, y_3, y_7, y_8) \\ y_4 \\ -\frac{1}{r}y_4 + f_2(y_1, y_3, y_7, y_8) \\ ry_1 \\ ry_3 \\ 0 \\ 0 \end{pmatrix}, \tag{55}$$

where $f_1$ and $f_2$ correspond to the right-hand sides of Eq. (53). This system is subject to the 8 boundary conditions specified at the center ($r = 0$) and outer edge ($r = L$):

$$\begin{gathered} y_2(0) = 0, \quad y_4(0) = 0, \quad y_5(0) = 0, \quad y_6(0) = 0, \\ y_2(L) = 0, \quad y_4(L) = 0, \quad y_5(L) = \frac{M_1}{2\pi}, \quad y_6(L) = \frac{M_2}{2\pi}. \end{gathered} \tag{56}$$

This boundary value problem is numerically integrated using a collocation method (implemented via MATLAB's bvp4c solver), yielding exact spatial profiles and equilibrium chemical potentials.

To relate the 2D simulations and 1D calculation, we use the circular domain of the 2D simulations where $r < L_0/2$ as the initial condition for the smallest $\bar{\phi}_2$ during the quasistatic addition of $\phi_2$ in the 1D calculation. As we quasistatically increase $\bar{\phi}_2$, we adopt the steady-state solution at each step as the initial guess for the next step.

After we obtain the equilibrium profile of the 1D problem with $L = L_0/2$, we apply a geometric correction to account for the outer-corner regions beyond the circular domain. Assuming the volume fractions in these corners equal the values at the outer boundary of the 1D calculation, $\phi_{2,\mathrm{dilute}} = y_3(L_0/2)$, the effective 2D average volume fraction $\bar{\phi}_{2,\mathrm{2D}}$ is evaluated as:

$$\bar{\phi}_{2,\mathrm{2D}} = \frac{1}{L_0^2}\left[2\pi y_6(L_0/2) + \phi_{2,\mathrm{dilute}}\left(L_0^2 - \frac{\pi L_0^2}{4}\right)\right], \tag{57}$$

where $y_6(L_0/2)$ is the cumulative 1D mass integrals for $\phi_2$. Details of the 1D BVP calculation are included in Table S1.

##### B3. Simulation Protocols

In this work, we mainly implement the following protocols of 2D numerical simulations:

1. **Adding $\phi_2$ to a solution with a preexisting condensate:** In this simulation, we initially simulate a binary system with a condensate enriched in $\phi_1$ at the center of the system. To accelerate this relaxation process, we prescribe the initial condensate morphology using a hyperbolic tangent profile. Once this two-component steady state is achieved, the client $\phi_2$ is added into the solvent. For rapid addition, the new solute $\phi_2$ replaces a portion of component $\phi_3$ at each spatial grid point $i$. Specifically, we set $\phi_{2i} = \phi_{3i}\bar{\phi}_2/\bar{\phi}_3$ to make sure $\bar{\phi}_2$ reaches the target value.

   We also implement a quasistatic addition, where $\phi_2$ increases at an infinitesimally slow rate, allowing the system to relax fully after each increment. In the quasistatic simulation, the first addition step is also rapid, but the system does not trigger secondary phase separation. In the following addition, each $\phi_2$ increment follows the same rule, i.e., $\Delta\phi_{2i} = \phi_{3i}\Delta\bar{\phi}_2/\bar{\phi}_3$.

2. **Simultaneous Rapid Addition:** In this simulation, both the scaffold and client are rapidly added simultaneously, i.e., $\phi_{2i} = \bar{\phi}_2$, $\phi_{1i} = \bar{\phi}_1$ for the initial condition for each lattice site.

For both the quasi-static and rapid addition scenarios, we add zero-mean and uniformly distributed random perturbations to the initial values with an amplitude of 1/20 of the average volume fraction of the added solute. During the simulations, we set an upper computational limit of $t = 5 \times 10^6$. The system state is evaluated every $\Delta t = 1000$, and the system is considered to have reached a steady state if the maximum change in the volume fractions of all components ($\phi_1$, $\phi_2$, and $\phi_3$) falls below $10^{-4}$.

During the simulations, if the maximum volume fraction of each of the three components exceeds 0.75 (note that the theoretical maximum volume fractions at equilibrium are $\phi_1 \approx 0.764$, $\phi_2 \approx 0.764$, and $\phi_3 \approx 0.920$) given the parameters we use in Fig. 3 and Fig. 4 of the main text), we consider the system has reached a three-phase state. We also use this criterion to detect the number of phases even if the simulation is terminated because it reaches the upper computational limit or reaches the steady state.

#### C. SUPPLEMENTARY FIGURES AND TABLES

TABLE S1. Parameters for the 1D boundary value problem (BVP).

| Parameter | Symbol | Value | Description |
|---|---|---|---|
| **I. Equation Parameters** | | | |
| Reduced Flory-Huggins interaction | $\chi_{11}, \chi_{22}$ | $-7$ | Self-interaction coefficients |
| | $\chi_{12}$ | Varies | Cross-interaction parameter ($\chi_{12} < 0$) |
| Dimensionless interfacial width | $\lambda$ | $4.5 \times 10^{-3}$ | Characteristic interface thickness |
| **II. Numerical Settings** | | | |
| 1D far-field boundary | $L$ | 0.5 | Coordinate of the outer computational boundary |
| BVP relative tolerance | RelTol | $10^{-4}$ | Relative error tolerance for the solver |
| BVP absolute tolerance | AbsTol | $10^{-6}$ | Absolute error tolerance for the solver |
| Initial mesh points | $N_{\text{init}}$ | 2000 | Number of grid points in the initial mesh |
| **III. Initial and Continuation Conditions** | | | |
| Scaffold average concentration | $\bar{\phi}_1$ | Varies | Global mean volume fraction of the scaffold |
| Client target concentration | $\bar{\phi}_2$ | Varies | Target mean volume fraction of the client |
| 1D BVP continuation step | $\Delta\bar{\phi}_2^{\text{1D}}$ | $10^{-4}$ | Step size for parameter continuation |

TABLE S2. Parameters for the 2D numerical simulations

.

| **Parameter** | **Symbol** | **Value** | **Description** |
|---|---|---|---|
| **I. Equation Parameters** | | | |
| Reduced Flory-Huggins interaction | $\chi_{11}, \chi_{22}$ | $-7$ | Self-interaction coefficients |
| | $\chi_{12}$ | Varies | Cross-interaction parameter ($\chi_{12} < 0$) |
| Dimensionless interfacial width | $\lambda$ | $4.5 \times 10^{-3}$ | Characteristic interface thickness |
| **II. Numerical Settings** | | | |
| Dimensionless system length | $L_0$ | 1.0 | Normalized length of the computational domain |
| Spatial grid resolution | $N_x \times N_y$ | $128 \times 128$, $256 \times 256$ | Case-dependent number of grid points |
| Spatial grid spacing | $\Delta x, \Delta y$ | 1/128, 1/256 | Spatial discretization step size |
| Dimensionless time step | $\Delta t$ | $10^{-1}$–$10^{1}$ | Constant time step (simulation-dependent) |
| Stabilization parameter | $A$ | $0.5 \max(\lvert\chi_{ij}\rvert)$ | Linear stabilization constant |
| **III. Initial Conditions** | | | |
| Scaffold average concentration | $\bar{\phi}_1$ | 0.1 (default) | Global mean volume fraction of the scaffold |
| Client target concentration | $\bar{\phi}_2$ | Varies | Target mean volume fraction of the client |
| 2D quasi-static addition step | $\Delta\bar{\phi}_2^{\mathrm{2D}}$ | $10^{-2}$ | Client increment per relaxation step |
| Initial noise amplitude | $\delta\phi$ | $\pm\bar{\phi}_1/20$ | Amplitude of random noise for initialization |

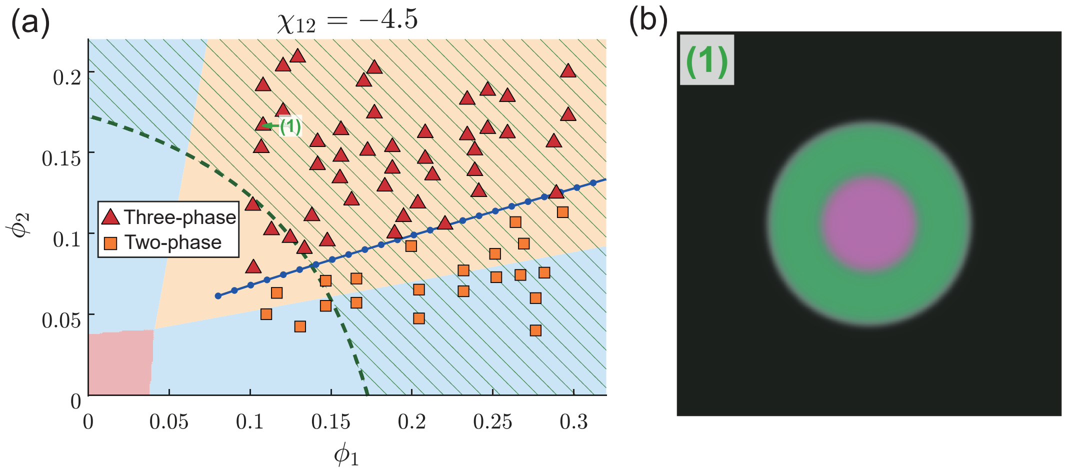


FIG. S1. Emergence of core-shell condensate morphologies under rapid $\phi_2$ addition. (a) Phase diagram identical to Fig. 3a, but highlighting a point $(\bar{\phi}_1, \bar{\phi}_2) = (0.108, 0.167)$ in the high-$\bar{\phi}_2$ regime. (b) Snapshot at $t = 5 \times 10^6$ corresponding to the marked parameter in panel a. In this simulation, the newly formed $\phi_2$ condensate completely encloses the primary $\phi_1$ condensate. As demonstrated in Fig. 2c of the main text, the absolute value of $\partial\mu_2/\partial\bar{\phi}_2$ decreases as $\bar{\phi}_2$ increases so that the gradient cost can suppress the interfacial instability, explaining why the core-shell morphology remains stable in this regime.

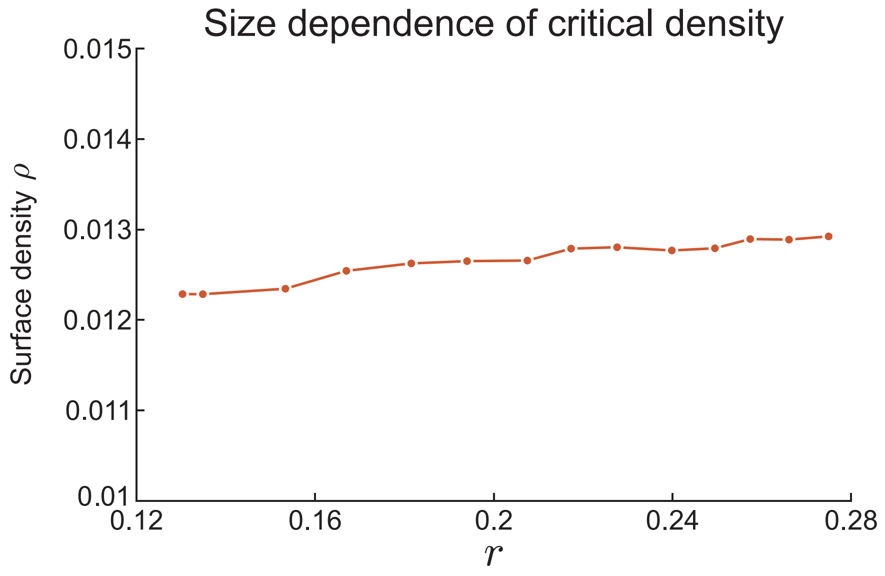


FIG. S2. Relationship between condensate radius and the critical surface density at interfacial instability. The horizontal axis represents the radius of the scaffold condensate($\phi_1$), and the vertical axis denotes the critical interfacial surface density ($\rho$) of the client component at the onset of phase separation. As the condensate radius increases, the critical surface density $\rho$ exhibits only a marginal increase. Therefore, in Fig. 4e of the main text, we set the value of the red dashed line as the average of the values shown here.

[1] M. Doi, *Soft matter physics* (Oxford University Press, 2013).

[2] U. M. Ascher, S. J. Ruuth, and R. J. Spiteri, Implicit-explicit runge-kutta methods for time-dependent partial differential equations, Applied Numerical Mathematics **25**, 151 (1997).

[3] J. Zhu, L.-Q. Chen, J. Shen, and V. Tikare, Coarsening kinetics from a variable-mobility cahn-hilliard equation: Application of a semi-implicit fourier spectral method, Physical Review E **60**, 3564 (1999).

[4] S. Mao, D. Kuldinow, M. P. Haataja, and A. Košmrlj, Phase behavior and morphology of multicomponent liquid mixtures, Soft Matter **15**, 1297 (2019).
